# rsfMRI-based Brain Entropy is negatively correlated with Gray Matter Volume and Surface Area

**DOI:** 10.1101/2024.04.28.591371

**Authors:** G. Del Mauro, Z. Wang

## Abstract

In recent years, brain entropy (BEN) has been ossociated with a number of neurocognitive, biological, and sociodemographic variables. However, its link with brain morphology is still unknown. In this study, we use resting-state fMRI (rsfMRI) data to estimate BEN maps and investigate their associations with three metrics of brain morphology: gray matter volume (GMV), surface area (SA), and cortical thickness (CT). Separate analyses will be performed on BEN maps derived from four distinct rsfMRI runs, and using both a voxelwise and a regions of interest (ROIs) approach. Our findings consistently showed that lower BEN (i.e., higher temporal coherence of brain activity) was related to increased GMV and SA in the lateral frontal and temporal lobes, inferior parietal lobules, and precuneus. We hypothesize that lower BEN and higher SA might both reflect higher brain reserve as well as increased information processing capacity.

## Introduction

In recent years, brain entropy (BEN) mapping has been increasingly adopted to characterize in a unique fashion the temporal aspect of human brain activity as measured by neuroimaging techniques, in particular resting-state fMRI (rsfMRI) (1). BEN provides an index of the “randomness” or “irregularity” of the brain activity. BEN estimated from rsfMRI using algorithms such as the Sample Entropy (SampEn) (2) refers to the repetitiveness of the blood-oxygen level dependent (BOLD) signal and represents an indirect proxy of the long-range temporal correlations (LRTC) that characterize the neural activity (3). High BEN values indicate independence between the data, low number of repeated patterns, and randomness (i.e., low temporal coherence), while low BEN values indicate persistence, repetition, and predictive behavior (i.e., high temporal coherence) (2). rsfMRI-based BEN has been correlated to a number of cognitive (e.g., fluid intelligence) (4–7), biological (e.g., age, sex) (4), and sociodemographic (e.g., education) variables, and BEN alterations have been found in neurological and psychiatric disorders (8–11).

While the relationship between BEN and brain morphology remains unknown, a number of indirect evidence suggest seem to suggest the presence of such relationship. For example, BEN increase with age (4), and aging is well-known to be correlated to the deterioration of the brain structure (12). Similarly, BEN is inversely correlated to education (4), which is directly related to brain morphology and usually considered a robust proxy of cognitive reserve (13). Finally, prior evidence indicates that females have a higher BEN than males (4). Differences between males and females in brain’s structure and functioning have been extensively documented (14–18), and while the exact nature of these differences is still undetermined, it has been repeatedly and consistently reported that males have larger brain volume than females (14–18).

Based on these evidence, the aim of this study is to use data from the Human Connectome Project – Young Adult dataset (HCP-Y) (19) to investigate for the first time the potential link between BEN and brain structure, as indexed by the gray matter volume (GMV). In particular, we hypothesize the GMV will be negatively correlated to BEN. In addition, it is worth noting that GMV is a composite measure resulting from the product between surface area (SA) and cortical thickness (CT), which represent genetically-independent properties of the brain cortex (20) driven by distinct cellular mechanisms. Indeed, neurons of the cerebral cortex are organized in columns perpendicular to the cerebral surface (21). According to the radial unit hypothesis, cortical SA is driven by the number of columns, while CT is driven by the number of cells within a column (22). To investigate the unique contributions of these properties of the brain morphology, we will separately analyze the relationship of BEN with both SA and CT. We hypothesize that BEN will be negatively correlated to either both or only one of these metrics. Analyses will be performed on a voxelwise level as well as using regions of interest (ROIs) from a pre-defined cortical parcellation. Moreover, consistency and repeatability of the results will be tested by analyzing the BEN maps derived from four distinct rsfMRI runs.

## Materials and methods

### Data

rsfMRI, sociodemographic, and behavioral data were downloaded from the HCP-Y dataset (19). 989 healthy young-adult participants were included in the sample (mean age = 28.76 ± 3.71 years; mean education = 14.95 ± 1.77 years; males/females = 452/537) (see following paragraph for information about the exclusion criteria). The rsfMRI protocol of HCP-Y includes the acquisition of four rsfMRI runs collected over two days. Each day involved the acquisition of a resting-state session (Rest1 on day 1, Rest2 on day 2) composed of two runs. All runs lasted 15 minutes, included 1200 timepoints, and were acquired with eyes open and using the same multi-band sequence. In order to compensate for image distortions induced by the long scan time, readout direction was from left to right for the first run of each session (Rest1 LR and Rest2 LR), and from right to left for the second run of each session (Rest1 RL and Rest2 RL). Other acquisition parameters include: repetition time (TR) = 720ms, echo time (TE) = 33.1ms, resolution = 2×2×2mm^3^, number of slices = 72. Downloaded data were already preprocessed using the HCP preprocessing pipeline, which includes correction for spatial distortions, head motion, and B_0_ distortions; registration to T1-weighted structural images; normalization to MNI space; global intensity normalization; masking out non-brain voxel; high-pass temporal filtering; independent component analyses (ICA)-based artifact removal (see (23) for a detailed description of the HCP-Y preprocessing pipeline). Before BEN calculation, rsfMRI data were smoothed with a full-width-at-half-maximum (FWHM) filter of 2.5 mm.

### BEN mapping

BEN was calculated only for participants who completed all rsfMRI runs. BEN mapping was performed for each rsfMRI run with the BEN mapping toolbox (BENtbx) (2). Briefly, BEN values were calculated using the SampEn formula, which is the logarithmic likelihood that a small section (within a time window of length ‘m’) of the data that “matches” with other sections will still “match” the others if the section window length increases by 1 (further information about BEN calculation procedure are reported in (2). “Match” is defined by a threshold of r times standard deviation of the entire time series. Window length m is widely set to be from 2 to 3. The embedding vector matching cut-off should be selected to avoid “no matching” (when it is too small) and “all matching” (when it is too big) (24). Both parameters have been assessed in previous publications (2). In this study, a window length of 3 and a cut-off threshold of 0.6 were adopted (2).

Participants may have altered BEN as a consequence of excessive head motions and artifacts at different time points, incorrect preprocessing, or abnormal brain state according to previous disease population studies (8). After entropy calculation, potential outliers were identified by first calculating, separately for each run, the mean BEN of each subject. Then, a participant was excluded from the subsequent analyses if the mean BEN of at least one run was above |3| standard deviations from the group average (5). A total of XX participants were identified as outliers and excluded.

Finally, average BEN values were extracted from 68 regions of the Desikan-Killiany cortical parcellation (25).

### Brain morphology metrics

The following summary metrics of the brain morphology were obtained using FreeSurfer (http://surfer.nmr.mgh.harvard.edu/): total GMV (mm^3^), total SA (mm^2^), and average CT (mm). In addition, GMV, SA, and CT values were extracted from the regions of the Desikan-Killiany cortical parcellation (25).

### Statistical analysis

#### Whole brain analyses

Whole brain analyses were performed using the Nilearn library running of Python 3 (26). Independently for each run (i.e., Rest1 LR, Rest1 RL, Rest2 LR, and Rest2 RL), three multiple regression models were estimated. In all models, the BEN maps of the participants were included as dependent variable, and age and sex as covariates. In addition, total GMV, total SA, and average CT were alternatively included as regressor. Then, the effect of each of these metrics was tested independently. Results were corrected for multiple comparisons at the voxel level using the family-wise error (FWE) method and were considered significant if p_voxel_-FWE < 0.05 (Z-threshold = 5.15). Results were plotted using the plotting module of Nilearn.

#### Regions of interest (ROIs) analyses

Each region of the Desikan-Killiany atlas included: four average BEN values (one for each rsfMRI run), and three structural metrics (GMV, SA, and CT). For all ROIs, the effect of each structural metric was tested independently on all four average BEN values. Specifically, for each ROI, multiple regression models were estimated to test the relationship between the average BEN and the morphology of that area. All multiple regression models included the average BEN values of the ROI as dependent variable. A structural metric of that ROI (i.e., GMV, SA, or CT) was alternatively included in the model as regressor to test its effect on the average BEN. In addition, age, sex, and the appropriate summary metric of brain morphology (total GMV, total SA, or average CT) were included as covariates. These analyses were performed for each of the four runs separately. The multiple regression models were estimated using the statsmodels library running on Python 3. Results were corrected for multiple comparisons independently for each run and morphology index using the Bonferroni correction. Results were considered significant if p-Bonferroni < 0.05. Results were plotted using the fsbrain library running on R Studio (27).

## Results

### Whole brain analyses

The total GMV was related to reduced BEN in the bilateral lateral frontal cortex, including superior, middle, and inferior frontal gyri, inferior parietal lobule, including angular and supramarginal gyri, middle/inferior temporal gyri, and precuneus (p_voxel_-FWE < 0.05) (Figure 1). The effect of the total SA showed a pattern similar to that of the total GMV (p_voxel_-FWE < 0.05) (Figure 2).

**Figure 1.**
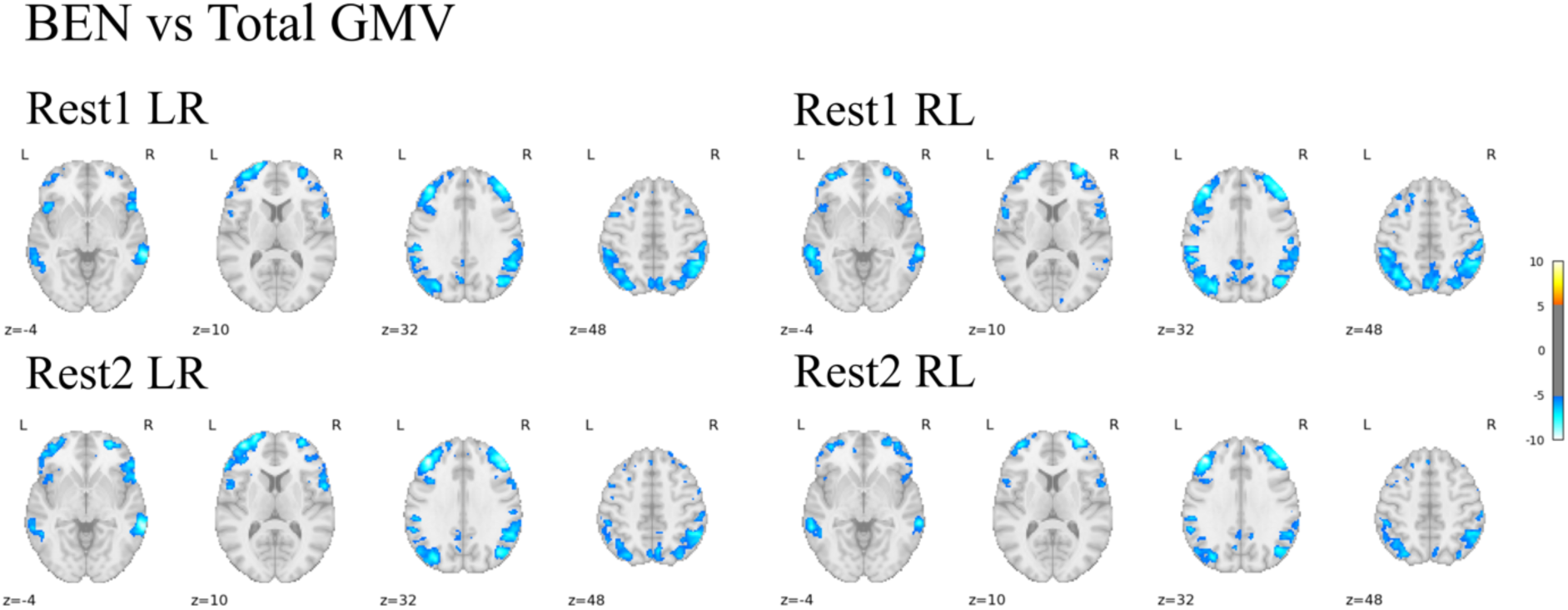
Effect of the total gray matter volume (GMV) on brain entropy (BEN). Age and sex were included as covariates. The effect of total GMV was tested on BEN maps derived from four distinct rsfMRI runs: Rest1 LR, Rest1 RL, Rest2 LR, Rest2 RL. Results were considered significant if p_voxel_-FWE < 0.05. Colorbar is based on Z-scores. The figure indicate only a negative relationship between BEN and total GMV, meaning that higher total GMV is related to lower BEN (cold colors).

**Figure 2.**
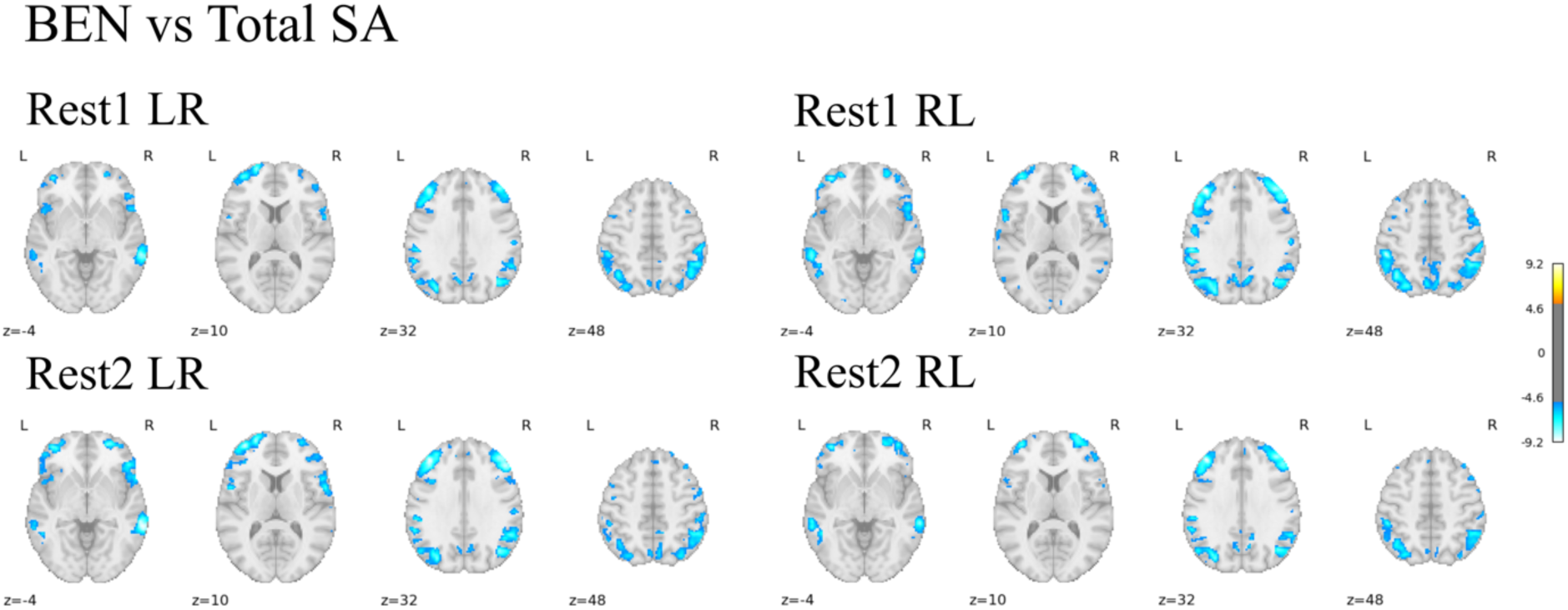
Effect of the total surface area (SA) on brain entropy (BEN). Age and sex were included as covariates. The effect of total SA was tested on BEN maps derived from four distinct rsfMRI runs: Rest1 LR, Rest1 RL, Rest2 LR, Rest2 RL. Results were considered significant if p_voxel_-FWE < 0.05. Colorbar is based on Z-scores. The figure indicate only a negative relationship between BEN and total SA, meaning that higher total SA is related to lower BEN (cold colors).

When the total GMV and the total SA were included in the model, the effect of sex was significant only in few small clusters (Figure S1, S2).

No significant effect of the average CT was observed. When the CT was included in the model, females showed overall higher BEN than males (Figure S3).

### Regional analyses

GMV, corrected for the total GMV, was significantly and negatively correlated to BEN in the lateral and medial frontal lobes, inferior parietal lobules, cingulate cortex, precuneus, and right middle temporal gyrus (p-Bonferroni < 0.05) (Figure 3; see Table 1 for labels of all significant areas and corresponding t-values).

**Figure 3.**
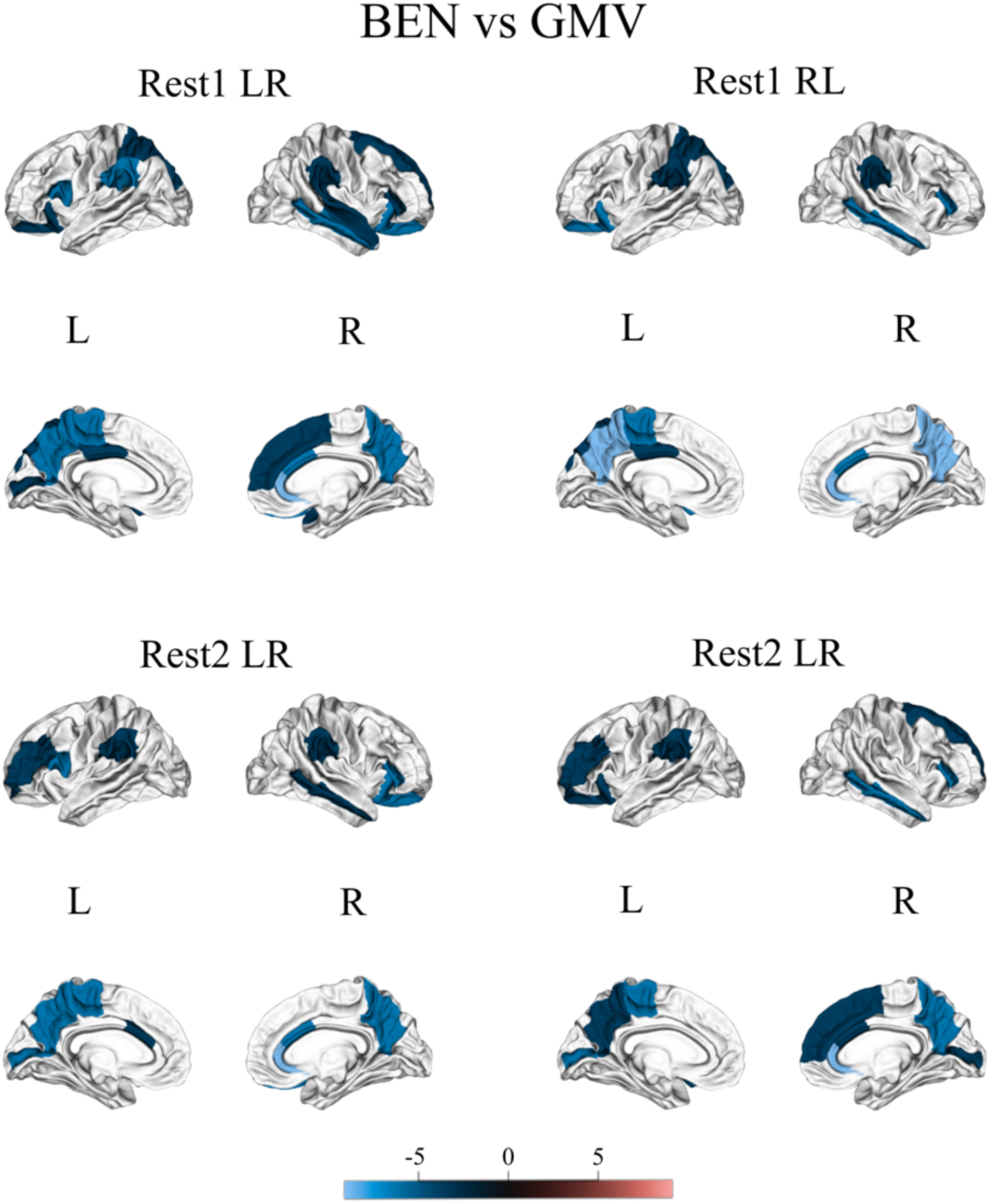
Regional analyses based on the Desikan-Killian cortical parcellation. For each area, the relationship between average brain entropy (BEN) and gray matter volume (GMV) was tested. Age, sex, and total GMV were included as covariates. Results were corrected for multiple comparisons using the Bonferroni correction and were considered significant if p-Bonferroni < 0.05. Colorbar is based on T-values. The figure indicate only negative relationships between local BEN and GMV (cold colors).

**Table 1.** Regions of Desikan-Killiany cortical parcellation showing a significant association between average brain entropy (BEN) and gray matter volume (GMV) in each of the four rsfMRI runs. For each region, the corresponding T-value is reported.

| GMV |  |  |
| --- | --- | --- |
| Run | Label | T-value |
| Rest1 LR | L Caudal anterior cingulate | -3.39 |
|  | L Paracentral | -4.99 |
|  | L Pars opercularis | -4.36 |
|  | L Pericalcarine | -4.64 |
|  | L Precuneus | -4.79 |
|  | L Rostral middle frontal | -3.62 |
|  | L Supramarginal | -3.99 |
|  | R Caudal anterior cingulate | -4.49 |
|  | R Lateral orbitofrontal | -4.59 |
|  | R Middle temporal | -3.89 |
|  | R Pars triangularis | -4.47 |
|  | R Precuneus | -4.93 |
|  | R Rostral anterior cingulate | -7.7 |
|  | R Supramarginal | -3.42 |
| Rest1 RL | L Lateral orbitofrontal | -4.59 |
|  | L Paracentral | -4.92 |
|  | L Posterior cingulate | -3.4 |
|  | L Precuneus | -6.31 |
|  | L Superior parietal | -3.51 |
|  | L Supramarginal | -3.42 |
|  | R Caudal anterior cingulate | -4.02 |
|  | R Middle temporal | -4.27 |
|  | R Pars triangularis | -4.68 |
|  | R Precuneus | -6.18 |
|  | R Rostral anterior cingulate | -6.35 |
|  | R Supramarginal | -3.85 |
| Rest2 LR | L Lateral orbitofrontal | -3.56 |
|  | L Paracentral | -5.24 |
|  | L Pars opercularis | -4.68 |
|  | L Pericalcarine | -3.56 |
|  | L Posterior cingulate | -3.4 |
|  | L Precuneus | -4.88 |
|  | L Superior frontal | -3.64 |
|  | L Superior parietal | -3.64 |
|  | L Supramarginal | -4.26 |
|  | R Caudal anterior cingulate | -4.26 |
|  | R Lateral orbitofrontal | -4.34 |
|  | R Middle temporal | -5.11 |
|  | R Pars triangularis | -3.93 |
|  | R Precuneus | -5.77 |
|  | R Rostral anterior cingulate | -6.28 |
|  | R Superior frontal | -3.91 |
|  | R Superior temporal | -3.43 |
|  | R Supramarginal | -3.84 |
|  | R Transverse temporal | -3.55 |
| Rest2 RL | L Lateral orbitofrontal | -3.6 |
|  | L Paracentral | -4.04 |
|  | L Pericalcarine | -5.14 |
|  | L Precuneus | -3.82 |
|  | L Rostral middlefrontal | -3.46 |
|  | L Supramarginal | -3.7 |
|  | R Caudal anterior cingulate | -3.93 |
|  | R Middle temporal | -4.95 |
|  | R Pars triangularis | -5.36 |
|  | R Pericalcarine | -3.62 |
|  | R Precuneus | -4.95 |
|  | R Rostral anterior cingulate | -7.04 |
|  | R Superior frontal | -3.78 |

A similar pattern was also observed when testing the effect of SA corrected for total SA (Figure 4, Table 2).

**Figure 4.**
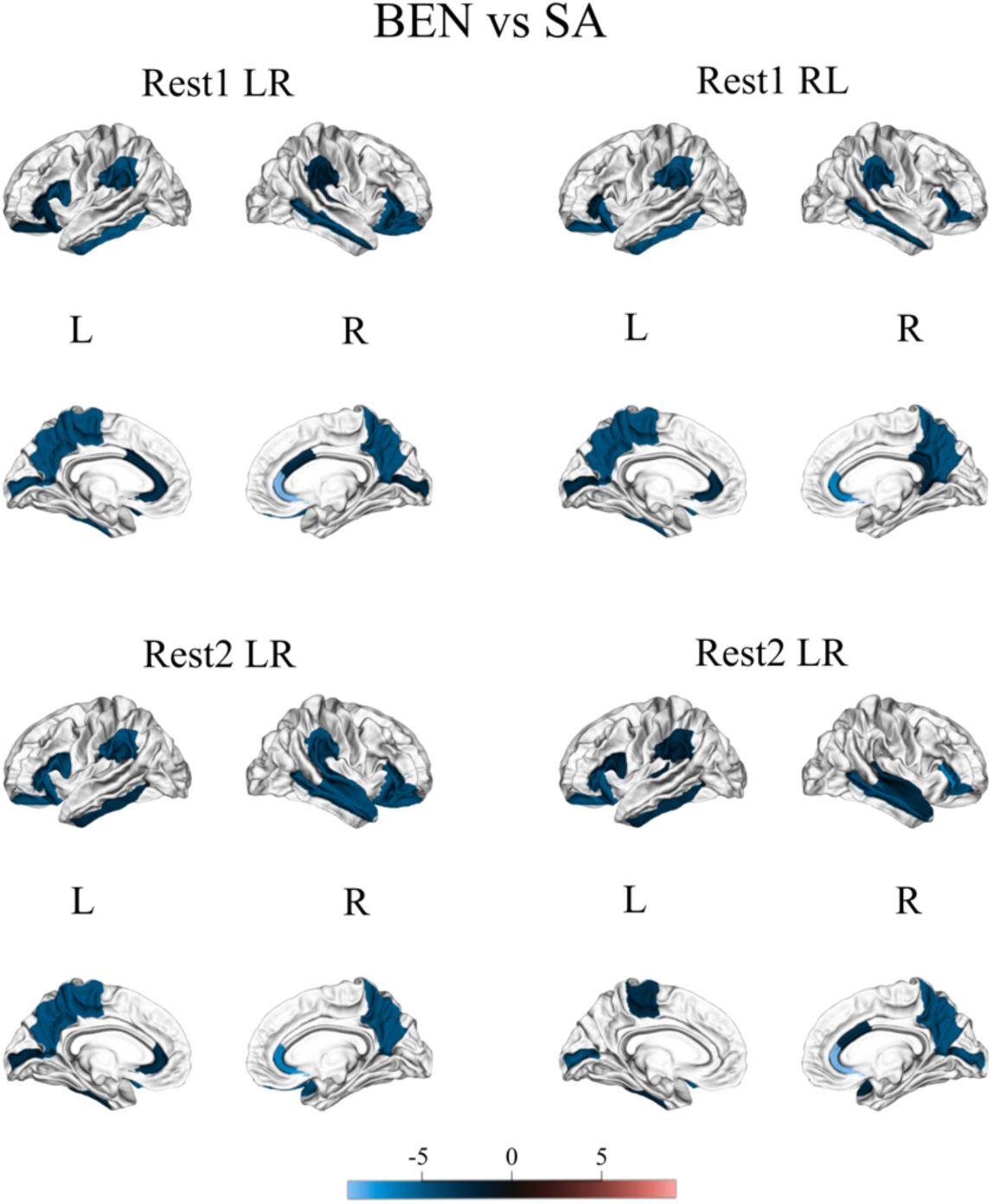
Regional analyses based on the Desikan-Killian cortical parcellation. For each area, the relationship between average brain entropy (BEN) and surface area (SA) was tested. Age, sex, and total SA were included as covariates. Results were corrected for multiple comparisons using the Bonferroni correction and were considered significant if p-Bonferroni < 0.05. Colorbar is based on T-values. The figure indicate only negative relationships between local BEN and SA (cold colors).

**Table 2.**
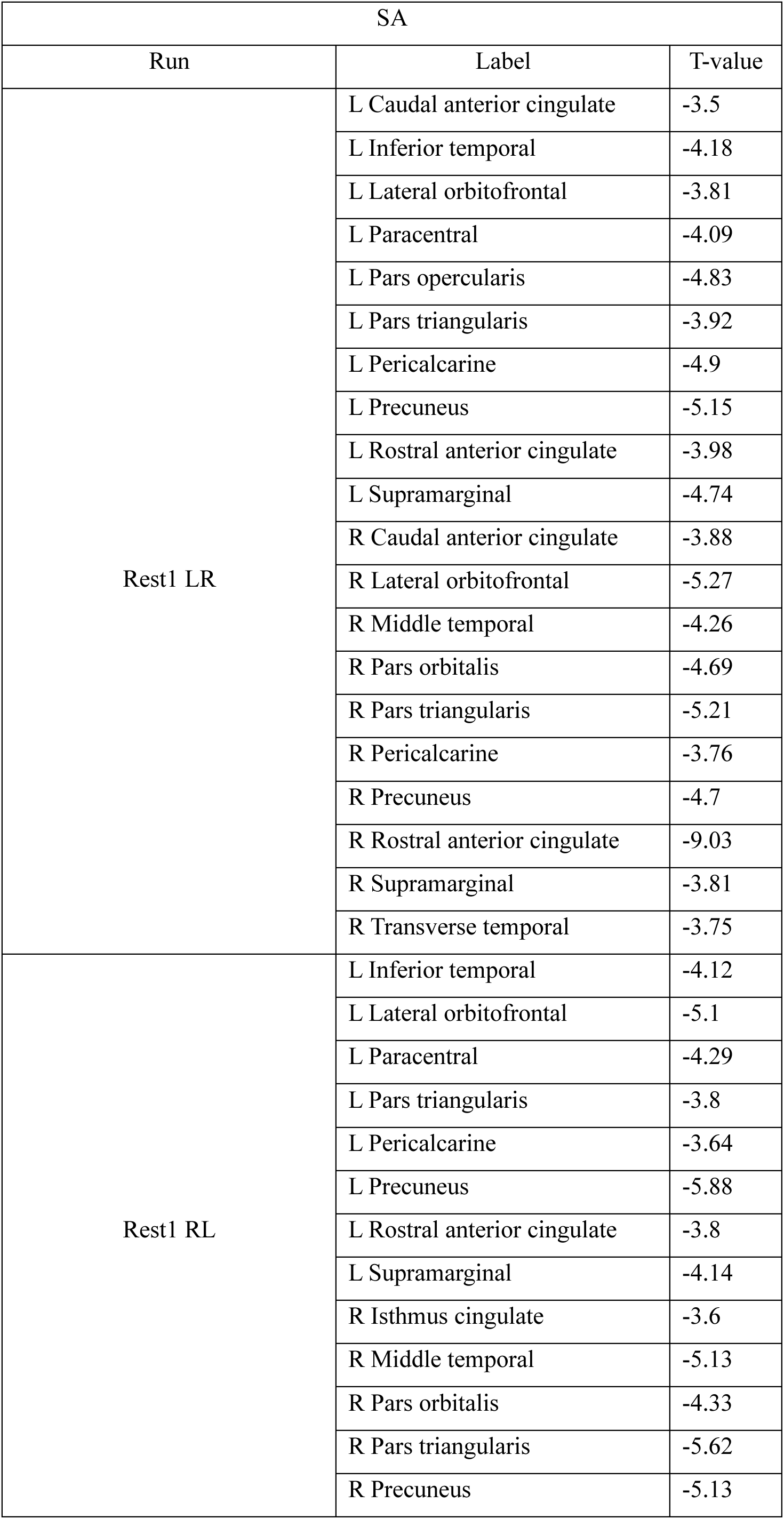
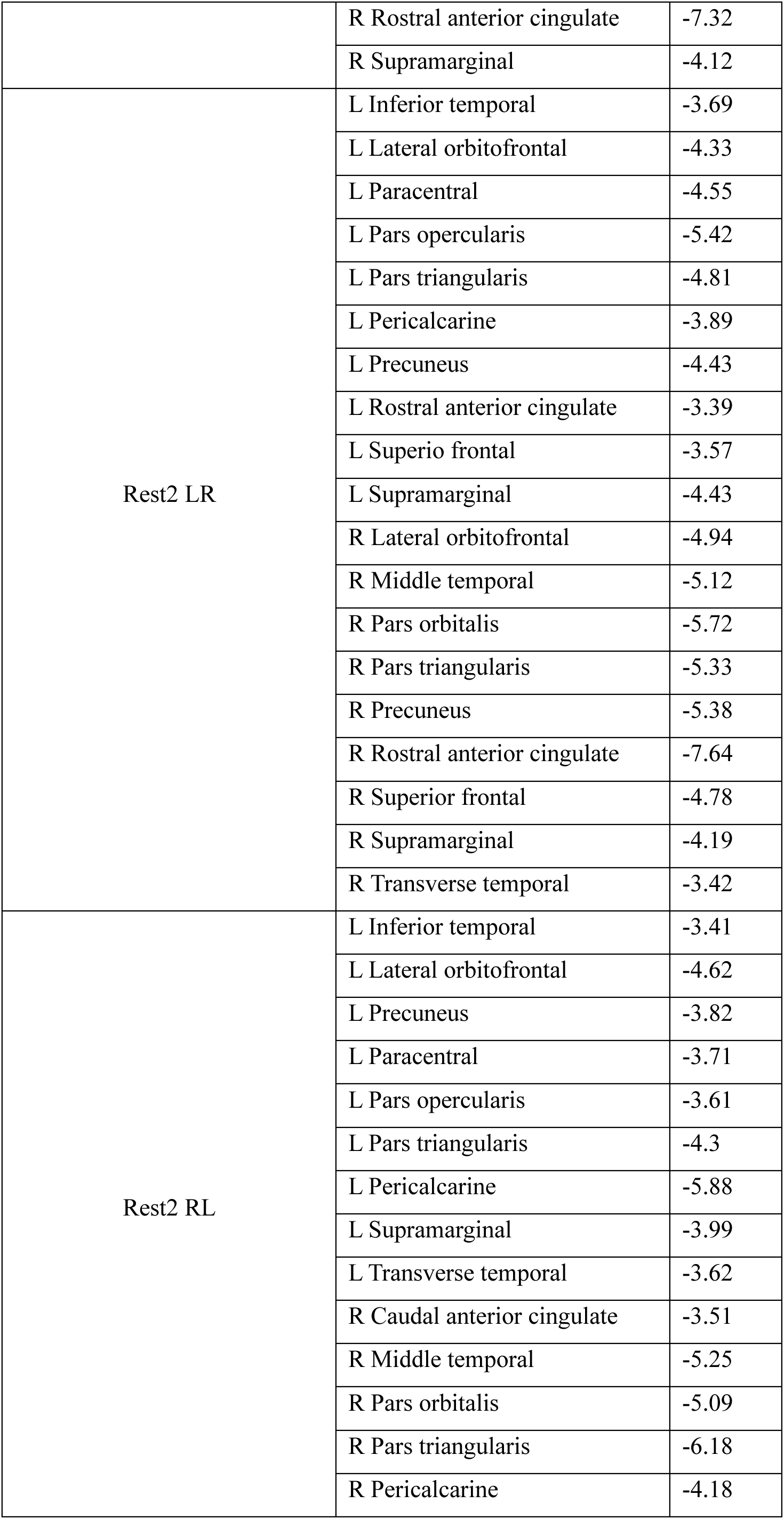
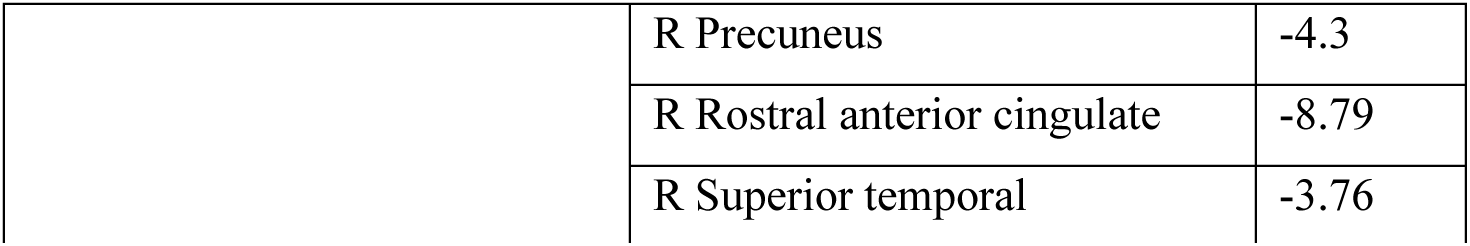
Regions of Desikan-Killiany cortical parcellation showing a significant association between average brain entropy (BEN) and surface area (SA) in each of the four rsfMRI runs. For each region, the corresponding T-value is reported.

| SA |  |  |
| --- | --- | --- |
| Run | Label | T-value |
| Rest1 LR | L Caudal anterior cingulate | -3.5 |
|  | L Inferior temporal | -4.18 |
|  | L Lateral orbitofrontal | -3.81 |
|  | L Paracentral | -4.09 |
|  | L Pars opercularis | -4.83 |
|  | L Pars triangularis | -3.92 |
|  | L Pericalcarine | -4.9 |
|  | L Precuneus | -5.15 |
|  | L Rostral anterior cingulate | -3.98 |
|  | L Supramarginal | -4.74 |
|  | R Caudal anterior cingulate | -3.88 |
|  | R Lateral orbitofrontal | -5.27 |
|  | R Middle temporal | -4.26 |
|  | R Pars orbitalis | -4.69 |
|  | R Pars triangularis | -5.21 |
|  | R Pericalcarine | -3.76 |
|  | R Precuneus | -4.7 |
|  | R Rostral anterior cingulate | -9.03 |
|  | R Supramarginal | -3.81 |
|  | R Transverse temporal | -3.75 |
| Rest1 RL | L Inferior temporal | -4.12 |
|  | L Lateral orbitofrontal | -5.1 |
|  | L Paracentral | -4.29 |
|  | L Pars triangularis | -3.8 |
|  | L Pericalcarine | -3.64 |
|  | L Precuneus | -5.88 |
|  | L Rostral anterior cingulate | -3.8 |
|  | L Supramarginal | -4.14 |
|  | R Isthmus cingulate | -3.6 |
|  | R Middle temporal | -5.13 |
|  | R Pars orbitalis | -4.33 |
|  | R Pars triangularis | -5.62 |
|  | R Precuneus | -5.13 |
|  | R Rostral anterior cingulate | -7.32 |
|  | R Supramarginal | -4.12 |
| Rest2 LR | L Inferior temporal | -3.69 |
|  | L Lateral orbitofrontal | -4.33 |
|  | L Paracentral | -4.55 |
|  | L Pars opercularis | -5.42 |
|  | L Pars triangularis | -4.81 |
|  | L Pericalcarine | -3.89 |
|  | L Precuneus | -4.43 |
|  | L Rostral anterior cingulate | -3.39 |
|  | L Superior frontal | -3.57 |
|  | L Supramarginal | -4.43 |
|  | R Lateral orbitofrontal | -4.94 |
|  | R Middle temporal | -5.12 |
|  | R Pars orbitalis | -5.72 |
|  | R Pars triangularis | -5.33 |
|  | R Precuneus | -5.38 |
|  | R Rostral anterior cingulate | -7.64 |
|  | R Superior frontal | -4.78 |
|  | R Supramarginal | -4.19 |
|  | R Transverse temporal | -3.42 |
| Rest2 RL | L Inferior temporal | -3.41 |
|  | L Lateral orbitofrontal | -4.62 |
|  | L Precuneus | -3.82 |
|  | L Paracentral | -3.71 |
|  | L Pars opercularis | -3.61 |
|  | L Pars triangularis | -4.3 |
|  | L Pericalcarine | -5.88 |
|  | L Supramarginal | -3.99 |
|  | L Transverse temporal | -3.62 |
|  | R Caudal anterior cingulate | -3.51 |
|  | R Middle temporal | -5.25 |
|  | R Pars orbitalis | -5.09 |
|  | R Pars triangularis | -6.18 |
|  | R Pericalcarine | -4.18 |
|  | R Precuneus | -4.3 |
|  | R Rostral anterior cingulate | -8.79 |
|  | R Superior temporal | -3.76 |

Finally, CT (corrected for the average CT) was negatively associated to BEN in the left paracentral gyrus, and positively associated to BEN in the left postcentral gyrus, and bilateral pars triangularis and orbitalis (Figure 5, Table 3).

**Figure 5.**
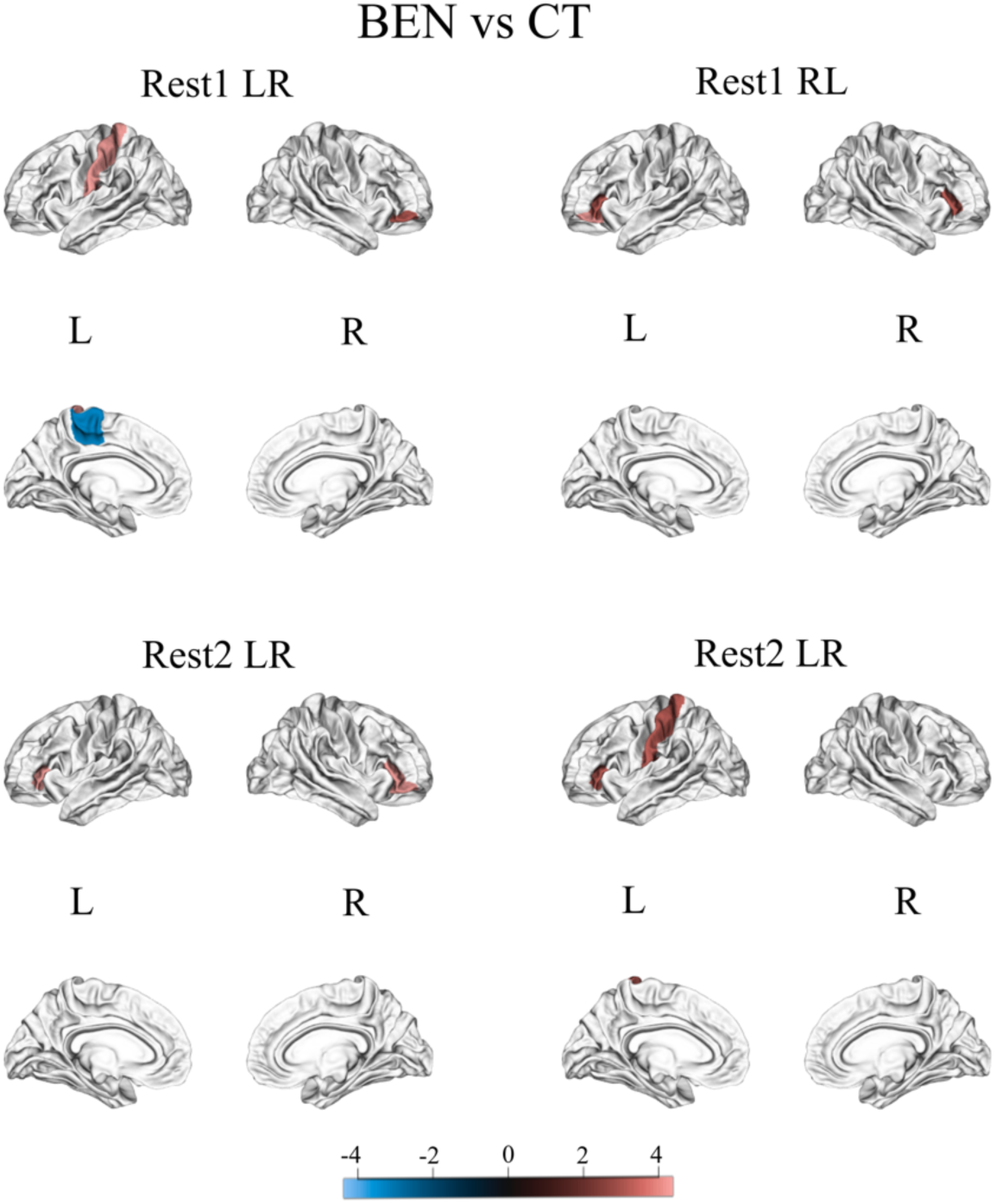
Regional analyses based on the Desikan-Killian cortical parcellation. For each area, the relationship between average brain entropy (BEN) and cortical thickness (CT) was tested. Age, sex, and average CT were included as covariates. Results were corrected for multiple comparisons using the Bonferroni correction and were considered significant if p-Bonferroni < 0.05. Colorbar is based on T-values. The figure indicate both positive (hot colors) and negative (cold colors) relationships between local BEN and CT.

**Table 3.** Regions of Desikan-Killiany cortical parcellation showing a significant association between average brain entropy (BEN) and cortical thickness (CT) in each of the four rsfMRI runs. For each region, the corresponding T-value is reported.

| CT |  |  |
| --- | --- | --- |
| Run | Label | T-value |
| Rest1 LR | L Paracentral | -3.63 |
|  | L Postcentral | 4.07 |
|  | R Pars orbitalis | 3.89 |
| Rest1 RL | L Pars triangularis | 3.71 |
|  | L Pars orbitalis | 4.33 |
|  | R Pars triangularis | 3.41 |
| Rest2 LR | L Pars triangularis | 4.02 |
|  | R Pars orbitalis | 4.03 |
|  | R Pars triangularis | 4.31 |
| Rest2 RL | L Pars triangularis | 3.53 |
|  | L Postcentral | 3.74 |

## Discussion

In this study, we investigated for the first time the relationship between BEN and brain morphology. Consistently with our hypothesis, GMV was inversely related to BEN, and additional analyses revealed that this effect is attributable to the SA rather than CT. Voxelwise and ROIs analyses both showed that this relationship is centered in the lateral frontal and temporal cortices, inferior parietal lobules as well as precuneus. ROIs analyses also displayed additional significant effects in the cingulate cortex and medial parietal lobes. Interestingly, when testing the effects of total GMV and SA, the BEN difference between males and females was negligible compared to when testing the effect of average CT as well as previous findings where females displayed higher BEN than males brain wide (4). This finding suggests that the BEN difference between males and females might be due to females having lower brain size than males (14, 15).

It was worth noting that BEN has been previously negatively correlated to general intelligence (5), and specifically fluid intelligence (4), in brain regions including lateral frontal and temporal cortices, inferior parietal lobules, and precuneus. In this work, we showed that the BEN of the same regions is also inversely related to the SA. A number of prior findings suggest that general cognitive ability might be correlated to SA rather than CT (28–31). For instance, it has been shown that the genetic association between GMV and general intelligence, as well as that the relationship between regional GMV and cognitive aging differences, are predominantly driven by SA (29, 31). Similarly, while CT has been found to be more correlated to aging, SA appears to be more related to fluid intelligence (28). Crucially, the pattern of regional correlations between SA and fluid intelligence is similar to the BEN-SA associations reported in this study (30).

Together, these findings indicate that BEN, and its relationship with neurocognition, might be closely related to the surface expansion of the cerebral cortex. Higher SA can lead to increased information processing capacity by increasing the number of cortical columns (32). Our results suggest the lower BEN, meaning higher temporal coherence of brain activity, is related to increased SA. Although the neural mechanisms underlying this relationship need to be investigated in future works, it is reasonable to hypothesize that both higher temporal coherence and SA reflect a capacity to more efficiently handle neural resourcers and process information. In addition, both BEN and SA might serve as proxies of the underlying brain reserve (33). Indeed, while the association between structural properties of the brain, including SA, and brain reserve has been widely documented (34–36), rsfMRI-derived BEN has also been recently proposed to reflect brain reserve, with lower BEN (i.e., higher temporal coherence) indicating a larger capacity of this reserve (4, 5).

The sample adopted in this study included only young adult participants. Since SA and CT are characterized by different developmental trajectories over the life span (37–39), future studies might investigate the relationship between BEN and brain morphology in a sample with a wider age-range. In addition, future works might explore whether the relationship between BEN and neurocognition is mediated by brain morphology, and in particular SA.

## Conclusions

This work investigated for the first time the relationship between BEN and brain structure using different indices of cortical morphology, including GMV, SA, and CT. Our results showed that lower BEN, corresponding to higher temporal coherence, is related to higher GMV and SA, but not CT. This relationship was observed in several brain regions including lateral frontal and temporal lobes, inferior parietal lobules, and precuneus. These results showed high consistency across different rsfMRI runs and analysis methods (i.e., voxelwise, ROIs). We hypothesize that higher functional temporal coherence and SA might reflect increased information process capacity as well as a higher brain reserve.

## Bibliography

1. Keshmiri S: Entropy and the Brain: An Overview. Entropy 2020; 22:917.

2. Wang Z, Li Y, Childress AR, Detre JA: Brain Entropy Mapping Using fMRI. PLoS One 2014; 9:e89948.

3. He BJ: Scale-Free Properties of the Functional Magnetic Resonance Imaging Signal during Rest and Task. The Journal of Neuroscience 2011; 31:13786–13795.

4. Wang Z: The neurocognitive correlates of brain entropy estimated by resting state fMRI. Neuroimage 2021; 232:117893.

5. Del Mauro G, Wang Z: Associations of Brain Entropy Estimated by Resting State fMRI With Physiological Indices, Body Mass Index, and Cognition. Journal of Magnetic Resonance Imaging 2023.

6. Camargo A, Del Mauro G, Wang Z: Task-induced changes in brain entropy. J Neurosci Res 2024; 102.

7. Shi L, Beaty RE, Chen Q, et al.: Brain Entropy is Associated with Divergent Thinking. Cerebral Cortex 2019.

8. Wang Z: Brain Entropy Mapping in Healthy Aging and Alzheimer’s Disease. Front Aging Neurosci 2020; 12.

9. Xue S-W, Yu Q, Guo Y, Song D, Wang Z: Resting-state brain entropy in schizophrenia. Compr Psychiatry 2019; 89:16–21.

10. Lin C, Lee S-H, Huang C-M, et al.: Increased brain entropy of resting-state fMRI mediates the relationship between depression severity and mental health-related quality of life in late-life depressed elderly. J Affect Disord 2019; 250:270–277.

11. Zhou F, Zhuang Y, Gong H, Zhan J, Grossman M, Wang Z: Resting State Brain Entropy Alterations in Relapsing Remitting Multiple Sclerosis. PLoS One 2016; 11:e0146080.

12. Zhao L, Matloff W, Ning K, Kim H, Dinov ID, Toga AW: Age-Related Differences in Brain Morphology and the Modifiers in Middle-Aged and Older Adults. Cerebral Cortex 2019; 29:4169–4193.

13. Boller B, Mellah S, Ducharme-Laliberté G, Belleville S: Relationships between years of education, regional grey matter volumes, and working memory-related brain activity in healthy older adults. Brain Imaging Behav 2017; 11:304–317.

14. Ruigrok ANV, Salimi-Khorshidi G, Lai MC, et al.: A meta-analysis of sex differences in human brain structure. Neurosci Biobehav Rev 2014; 39:34–50.

15. Ritchie SJ, Cox SR, Shen X, et al.: Sex differences in the adult human brain: Evidence from 5216 UK biobank participants. Cerebral Cortex 2018; 28:2959–2975.

16. Lotze M, Domin M, Gerlach FH, et al.: Novel findings from 2,838 Adult Brains on Sex Differences in Gray Matter Brain Volume. Sci Rep 2019; 9:1–7.

17. Decasien AR, Guma E, Liu S, Raznahan A: Sex differences in the human brain : a roadmap for more careful analysis and interpretation of a biological reality. Biol Sex Differ 2022:1–21.

18. Williams CM, Peyre H, Toro R, Ramus F: Neuroanatomical norms in the UK Biobank: The impact of allometric scaling, sex, and age. Hum Brain Mapp 2021; 42:4623–4642.

19. Van Essen DC, Smith SM, Barch DM, Behrens TEJ, Yacoub E, Ugurbil K: The WU-Minn Human Connectome Project: An overview. Neuroimage 2013; 80:62–79.

20. Panizzon MS, Fennema-Notestine C, Eyler LT, et al.: Distinct genetic influences on cortical surface area and cortical thickness. Cerebral Cortex 2009; 19:2728–2735.

21. Mountcastle V: The columnar organization of the neocortex. Brain 1997; 120:701–722.

22. Rakic P: Specification of Cerebral Cortical Areas. Science (1979) 1988; 241:170–176.

23. Glasser MF, Sotiropoulos SN, Wilson JA, et al.: The minimal preprocessing pipelines for the Human Connectome Project. Neuroimage 2013; 80:105–124.

24. Richman JS, Moorman JR: Physiological time-series analysis using approximate entropy and sample entropy. American Journal of Physiology-Heart and Circulatory Physiology 2000; 278:H2039–H2049.

25. Desikan RS, Ségonne F, Fischl B, et al.: An automated labeling system for subdividing the human cerebral cortex on MRI scans into gyral based regions of interest. Neuroimage 2006; 31:968–980.

26. Abraham A, Pedregosa F, Eickenberg M, et al.: Machine learning for neuroimaging with scikit-learn. Front Neuroinform 2014; 8.

27. Schaefer T, Ecker C: fsbrain: an R package for the visualization of structural neuroimaging data. bioRxiv 2020.

28. Borgeest G, Henson R, Kietzmann T, et al.: A morphometric double dissociation: cortical thickness is more related to aging; surface area is more related to cognition. bioRxiv 2021.

29. Cox SR, Bastin ME, Ritchie SJ, et al.: Brain cortical characteristics of lifetime cognitive ageing. Brain Struct Funct 2018; 223:509–518.

30. Tadayon E, Pascual-Leone A, Santarnecchi E: Differential Contribution of Cortical Thickness, Surface Area, and Gyrification to Fluid and Crystallized Intelligence. Cerebral Cortex 2020; 30:215–225.

31. Vuoksimaa E, Panizzon MS, Chen CH, et al.: The Genetic Association between Neocortical Volume and General Cognitive Ability Is Driven by Global Surface Area Rather Than Thickness. Cerebral Cortex 2015; 25:2127–2137.

32. Rakic P: Confusing cortical columns. Proceedings of the National Academy of Sciences 2008; 105:12099–12100.

33. Stern Y: Cognitive reserve in ageing and Alzheimer’s disease. Lancet Neurol 2012; 11:1006–1012.

34. Seyedsalehi A, Warrier V, Bethlehem RAI, Perry BI, Burgess S, Murray GK: Educational attainment, structural brain reserve and Alzheimer’s disease: a Mendelian randomization analysis. Brain 2023; 146:2059–2074.

35. van Loenhoud AC, Groot C, Vogel JW, van der Flier WM, Ossenkoppele R: Is intracranial volume a suitable proxy for brain reserve? Alzheimers Res Ther 2018; 10:91.

36. Stern Y, Arenaza-Urquijo EM, Bartrés-Faz D, et al.: Whitepaper: Defining and investigating cognitive reserve, brain reserve, and brain maintenance. Alzheimer’s & Dementia 2020; 16:1305–1311.

37. Storsve AB, Fjell AM, Tamnes CK, et al.: Differential Longitudinal Changes in Cortical Thickness, Surface Area and Volume across the Adult Life Span: Regions of Accelerating and Decelerating Change. Journal of Neuroscience 2014; 34:8488–8498.

38. Wierenga LM, Langen M, Oranje B, Durston S: Unique developmental trajectories of cortical thickness and surface area. Neuroimage 2014; 87:120–126.

39. Sele S, Liem F, Mérillat S, Jäncke L: Age-related decline in the brain: a longitudinal study on inter-individual variability of cortical thickness, area, volume, and cognition. Neuroimage 2021; 240:118370.

