## Supplementary Figures for "rsfMRI-based Brain Entropy is negatively correlated with Gray Matter Volume and Surface Area"

Del Mauro G.[1], PhD

Wang Z.[1], PhD

[1]Department of Diagnostic Radiology and Nuclear Medicine, University of Maryland Schoold of Medicine

Address for correspondence:

Ze Wang, PhD, Professor

Department of Diagnostic Radiology & Nuclear Medicine

University of Maryland School of Medicine

670 W Baltimore St, HSF III, R1173

Baltimore 21202, MD

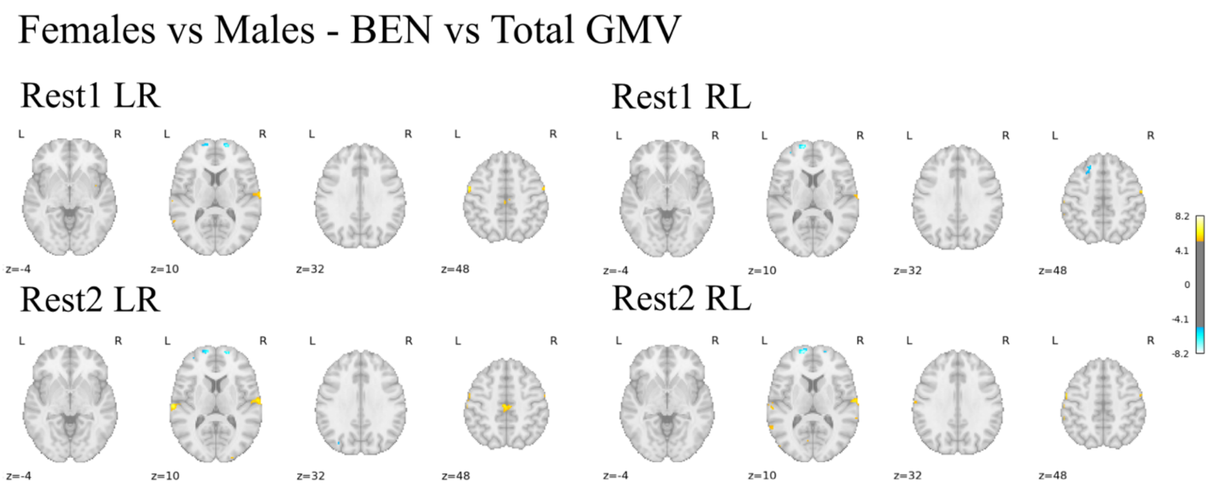


Figure S1. Brain Entropy (BEN) differences between males and females when including the total gray matter volume (GMV) as covariate in the multiple regression model. The effect of sex was tested on BEN maps derived from four distinct rsfMRI runs: Rest1 LR, Rest1 RL, Rest2 LR, Rest2 RL. Results were considered significant if $p_{voxel}$-FWE < 0.05. Colorbar is based on Z-scores. Warm colors indicate higher BEN in females, while cold colors indicate higher BEN in males.

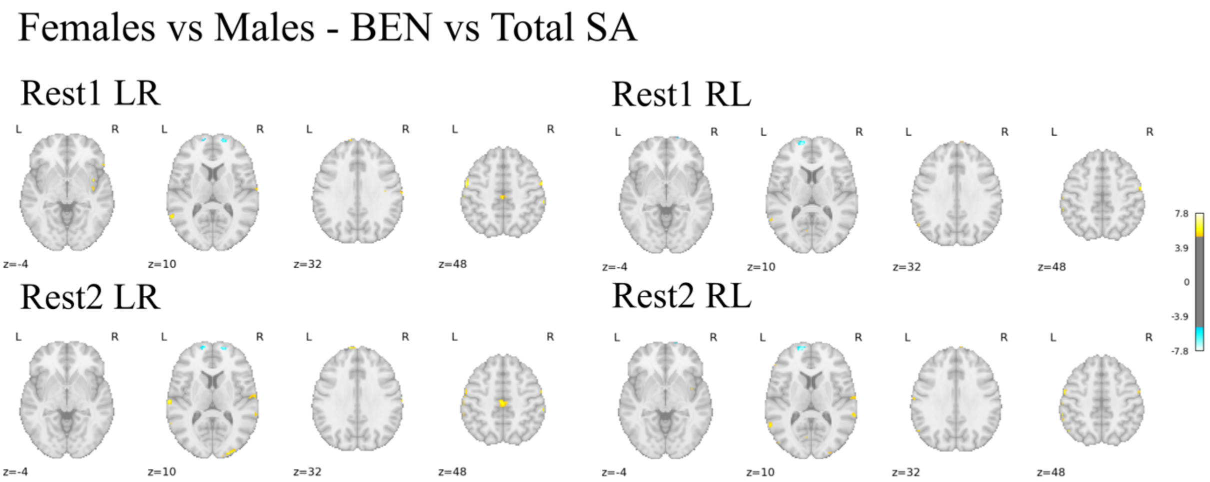


Figure S2. Brain Entropy (BEN) differences between males and females when including the total surface area (SA) as covariate in the multiple regression model. The effect of sex was tested on BEN maps derived from four distinct rsfMRI runs: Rest1 LR, Rest1 RL, Rest2 LR, Rest2 RL. Results were considered significant if $p_{voxel}$-FWE < 0.05. Colorbar is based on Z-scores. Warm colors indicate higher BEN in females, while cold colors indicate higher BEN in males.

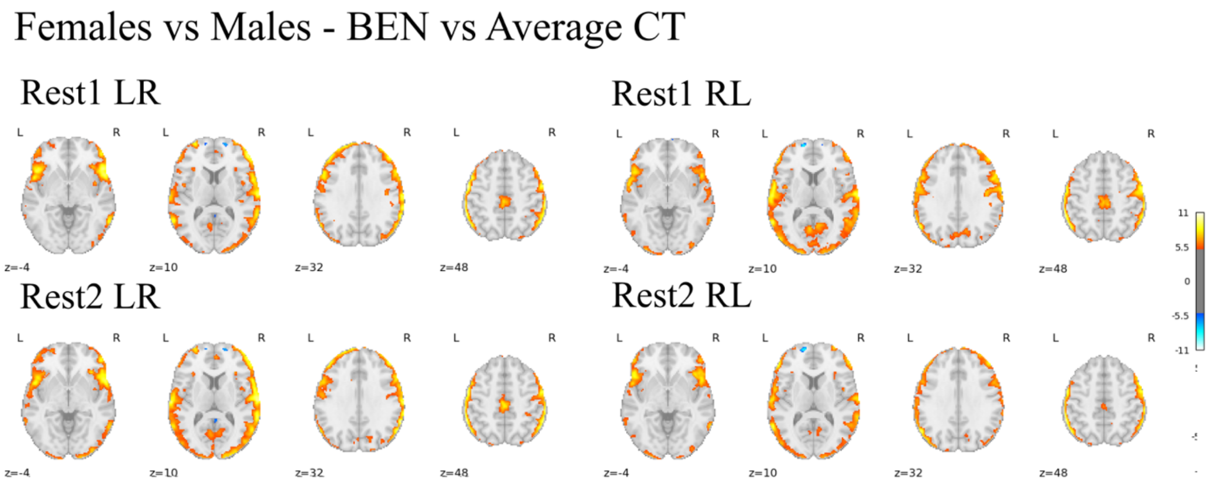


Figure S3. Brain Entropy (BEN) differences between males and females when including the average cortical thickness (CT) as covariate in the multiple regression model. The effect of sex was tested on BEN maps derived from four distinct rsfMRI runs: Rest1 LR, Rest1 RL, Rest2 LR, Rest2 RL. Results were considered significant if $p_{voxel}$-FWE < 0.05. Colorbar is based on Z-scores. Warm colors indicate higher BEN in females, while cold colors indicate higher BEN in males.
